# Orb-weaving spiders show a correlated syndrome of morphology and web structure in the wild

**DOI:** 10.1101/767335

**Authors:** David N. Fisher, Justin Yeager, Jonathan N. Pruitt

## Abstract

Extended phenotypes are traits that exist outside the physical body of the organism. Despite their potential role in the lives of both the organisms that express them and other organisms that can be influenced by extended phenotypes, the consistency and covariance with morphological and behaviour traits of extended phenotypes is rarely evaluated, especially in wild organisms. We repeatedly measured an extended phenotype that directly influences an organism’s prey acquisition, the web structure, of wild orb-weaving spiders (*Micrathena vigorsii*), which re-build their webs each day. We related web structure traits to behavioural traits and body size (length). Both web diameter and web density were repeatably different among individuals, while reaction to a predation threat was slightly so, but response to a prey stimulus and web symmetry were not. There was a syndrome between morphology and web structure traits, where larger spiders spun webs that were wider, had webs with increased thread spacing, and the spider tended to react more slowly to a predation threat. When a spider built a relatively larger web it was also relatively a less dense and less symmetrical web. The repeatability of web construction and relationship with spider body size we found may be common features of intra-population variation in web structure in spiders. Individual variation along the morphology and web structure syndrome could represent variation in individual foraging strategies, or age-based correlated changes. By estimating the consistency and covariances of extended phenotypes we can begin to evaluate what maintains their variation and how they might evolve.

## Introduction

Some phenotypes of organisms are “extended”, in that they exist beyond the physical body of the organism (Dawkins 1978; Dawkins 1982). Examples include bird nests, beaver dams, and spider webs. Extended phenotypes can relate to the survival, foraging, or mating success of an individual, and those that use the environment the individual modifies (including “ecosystem engineers”; Jones et al. 1994; Jones et al. 1997; Rosell et al. 2005; Kooch and Jalilvand 2008; Jones et al. 2009; White and O’Donnell 2010; Ransom 2011; Posthumus et al. 2015; Ringler et al. 2015; Fisher et al. 2019). When extended phenotypes are repeatably expressed over the life of an organism, they have the potential to show reversible plasticity, for example a bird may build a large nest one year and a small nest the following year. Despite this plasticity, traits with the potential to be labile are often consistent within an individual across time or in different environments, such that within a population there can be considerable variation among-individuals in the mean trait they express (Bell et al. 2009). Causes and consequences for this consistent among-individual variation has been the source of great interest, especially within the last 20 years (Koolhaas et al. 1999; Dall et al. 2004; Réale et al. 2007; Bell et al. 2009; Stamps and Groothuis 2010). However, plasticity and consistency of extended phenotypes is not extensively documented (Venner et al. 2000; Blamires 2010; Dirienzo and Montiglio 2016; Montiglio and DiRienzo 2016; Blamires, Hasemore, et al. 2017; Blamires, Martens, et al. 2017). Furthermore, the expression of extended phenotypes may covary with other repeatably expressed traits such as behaviours, or more stable straits such as morphology, but again this is not often documented (Dirienzo and Montiglio 2016; Montiglio and DiRienzo 2016). Yet, it is important that we do so as extended phenotypes alter the environment an individual experiences, possibly changing selection pressures both for itself and for other organisms that use the same environment.

Evaluating the repeatability of extended phenotypes and their associations with other aspects of behaviour and morphology has a variety of implications. First, as only a portion of any among-individual variation will have a genetic basis, repeatability can set the upper limit for heritability (Falconer 1981; Boake 1989; but see: Dohm 2002), and repeatability in behaviour has various consequences for ecological and evolutionary processes (Dall et al. 2012; Sih et al. 2012; Wolf and Weissing 2012). Meanwhile, extended phenotypes being correlated into “syndromes” with traits such as body size or boldness could help explain the maintenance of variation in extended phenotypes within populations (Roff 1992; Sih, Bell, Johnson, et al. 2004; Sih, Bell, and Johnson 2004). Further, trait associations can influence how syndrome structure (i.e., the G-matrix) and its constituent traits evolve (Lande 1979; Lande and Arnold 1983), which can sometimes impinge on adaptive evolution (Dochtermann and Dingemanse 2013; Royauté et al. 2019). The repeatability of extended phenotypes and their phenotypic integration with other traits is especially interesting because extended phenotypes help to engineer the external environment surrounding an organism, and therefore how selection acts on other traits. It is therefore necessary that we quantify the repeatability of extended phenotypes and assess how they are correlated with other traits.

Here we quantify the degree of consistency of various aspects of an extended phenotype and measure its covariance with body size and two behavioural traits. We studied the orb-weaving spider *Micrathena vigorsi* (Araneae: Araneidae; Fig. 1a). While spider webs were once thought to rigidly follow species specific patterns, more recent research has identified both among- and within-individual variation in web structure, such as differences in size, shape, the number of different kinds of threads/lines, and the number of mistakes made in the web’s construction (Sherman 1994; Heiling and Herberstein 2000; Venner et al. 2000; Blamires 2010; Dirienzo and Montiglio 2016; Montiglio and DiRienzo 2016). See Heiling and Herberstein (2000) for a review. Variation in web structure can influence what and how many prey are caught (Uetz et al. 1978; Chacon and Eberhard 1980; Sensenig et al. 2010), whether prey are retained (Blackledge and Zevenbergen 2006; Blamires, Martens, et al. 2017), how much protection the web provides from predators (Zevenbergen et al. 2008), and may also reflect a spider’s body condition (Blackledge and Zevenbergen 2007; Dirienzo and Montiglio 2016 Jul), recent experience (Nakata and Ushimaru 1999; Venner et al. 2000), and/or age (Anotaux et al. 2012; Anotaux et al. 2014). Spider web structure is therefore an important phenotype for various components of a spider’s fitness.

**Figure 1.**
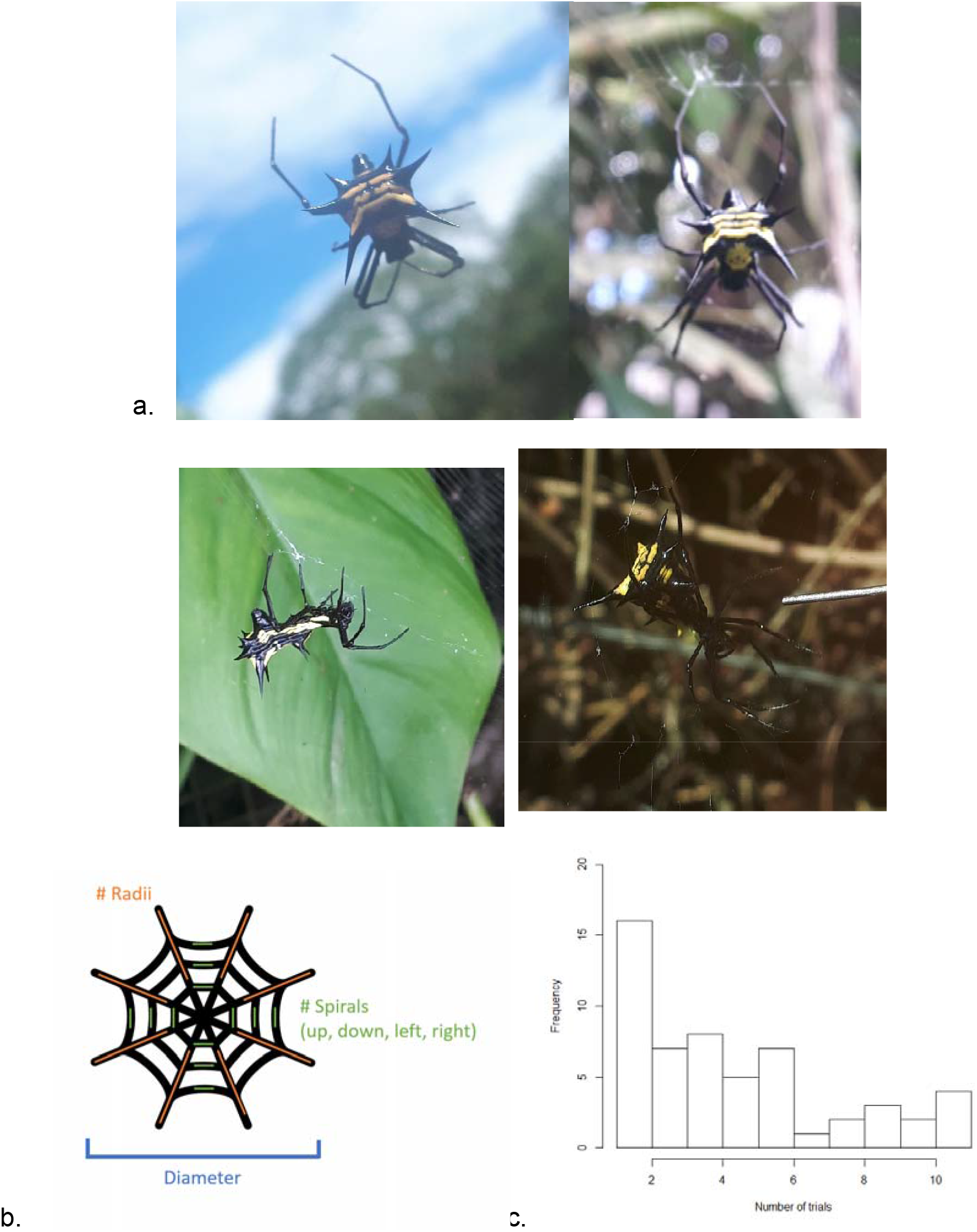
a. Pictures of *Micrathena vigorsii* (all DN Fisher). b. A diagram showing the different web characteristics used to calculate measures of web structure. c. Histogram of the number of times individuals were tested for web structure.

*Micrathena spp.* rebuild the orb of their web anew each day (Shelly 1984; Hodge 1987), and so allow one to easily measure the repeated expression of an extended phenotype and determine if individuals consistently differ from each other over time. It also allows us to see if there is day-to-day variation in web structure, if for example certain environmental conditions consistently influence web structure for all individuals. Previous studies on among-individual consistency in web structure have been conducted in captivity (Dirienzo and Montiglio 2016; Montiglio and DiRienzo 2016; Dirienzo and Montiglio 2016 Jul; DiRienzo and Dornhaus 2017; DiRienzo and Aonuma 2018), but our study took place in the wild to assess the degree of consistency of behaviour and web structure, and how they covary with each other and with spider body size, in a natural setting. Studies in standardised laboratory settings are useful when we want to control for external factors, but studies in free-living populations help us assess the degree of consistency and correlations between traits that natural selection may be acting upon. Conducting studies in the wild also avoids the potential confound that individual differences in phenotypes can be driven by individual differences in response to laboratory conditions (Roche et al. 2016).

We estimated the phenotypic correlations between spider predation and anti-predator behaviour, web structure traits, and body length. We then partitioned our estimated covariances into the among-individual, among-date, and residual levels. We predicted that larger spiders will build bigger and less dense webs (webs with fewer threads per area of the web), as they will be targeting larger prey (Uetz et al. 1978; Chacon and Eberhard 1980; Sensenig et al. 2010), and that larger spiders will more aggressively attack a prey stimulus and will react less readily towards a predator stimulus (Rundle and Brönmark 2001; Mayer et al. 2016). We also predicted that there will be a positive among-date covariance between responses to predator and prey stimuli, as on hotter days spiders might be more responsive in general to a range of different cues (Pruitt et al. 2011; Briffa et al. 2013). Finally, we predicted a negative residual covariance between web diameter and both web density and web symmetry, as on instances that a spider builds a relatively bigger web, it may also build a less dense and less symmetrical web (Blamires 2010). We do not expect a relationship here between web density and anti-predator behaviour (e.g. Blackledge and Zevenbergen 2007) as *M. vigorsii* do not use their web for defence. We investigated covariances among six traits, all at the among-individual, among-date, and residual level (with some exceptions, see below). This totals 45 covariances; we do not make predictions for each for the sake of brevity.

## Methods

### Data collection

Our study took place between 16/6/ – 15/7 2019 near Tena in Ecuador (approx. lat long = −1.044, −77.715), under the Ecuadorian Ministry of the Environment permit no. 014-2019-IC-FLO-DNB/MA. We located individual *M. vigorsii* (n = 55) in a hedgerow along a transect at the northern side of a road (route number: 436) between 9 and 121 cm above the ground (mean = 59 cm). We assume all individuals in our study are adult females, as males of *Micrathena* spp. are much smaller and stop building webs upon reaching maturity (Chickering 1961; Shelly 1984; Hodge 1987). Once we found an individual, we marked its location with a piece of flagging tape with the unique ID of the spider written on it. This allowed us to return and phenotype the same individual on different days without disrupting the individual’s behaviour by capturing it and marking it directly. While *Micrathena* spp. do rebuild their orb each day, the “frame” of the web remains in place, and so the position of the web will vary minimally day-to-day. When two individuals were close together, we used their size measurements and notes on individual characteristics such as colouration and exact web location to identify them. We are therefore confident that we were able to repeatably find and identify the same individuals. We only conducted testing between 14:00-17:00, after web construction should have been completed, limiting the impact the time of day could have on variation in behaviour and web structure.

The first time we found an individual, we measured its length (to the nearest 0.01 mm) using a pair of callipers (Traceable, Fisher Scientific, PA, USA) held up against the spider in its web. In 10/55 of cases (18.2%; mostly at the start of the study before the callipers were available) we did not measure the length of the individual. Following this, we tested the individual for responsiveness towards a prey stimulus. To do this, we touched a piece of wire attached to a modified vibrating device (8” Vibrating Jelly Dong, Top Cat Toys, Chatsworth CA, USA) to the lowest point of the orb of the web. Assays like this are commonly used to estimate foraging aggression in both solitary and social spiders (Laskowski and Pruitt 2014; Dirienzo and Montiglio 2016; Lichtenstein et al. 2019). We timed from the start of the vibrations until the spider touched the end of the vibrating wire. If the spider did not respond within 180 seconds (55/210 tests that the spider did not flee before the trial began) we set the spider’s score at 180. Next, we waited for the spider to return to the centre of the web and adopt a resting position (facing down towards the floor), before testing it for a reaction to a predation threat. We tapped the abdomen of the spider lightly with the extended lead of a mechanical pencil and recorded how many taps transpired until the spider fled. If the spider did not react within 24 taps (26/210 tests the spider did not flee before the trial began), we set the spider’s score at 24. These tests gave us measures of two behaviours: responsiveness to prey and reaction to a predator threat.

To quantify web structure, we counted the of number “radii” of the orb (strands extending from the centre outwards), and the number of “spirals” (strands perpendicular to the radii circling the centre) extending up, down, left and right from the centre of the web to its perimeter (giving four separate counts; Fig. 1b; Sensenig et al. 2010). We then measured the diameter of the web (to the nearest 1cm). We calculated web density as the number of radii multiplied by the mean number of spirals, divided by the web area (assuming a circular shape). Unfortunately, at the time of data collection we were not aware of Herberstein and Tso’s work, which indicates an ellipse is preferable is a circle for estimating web area (Herberstein and Tso 2000, although assuming a circle still gives reasonable approximations of true capture area; see also: Venner et al. 2001 for estimating the total capture thread length). We calculated web symmetry as the variance among the four counts of the number of spirals; for this measure lower values indicate a more symmetrical web (the correlation between this value and the coefficient of variation in spiral counts was 0.936). We also counted the number of lines attaching the web to the surrounding vegetation and measured the centre of the web’s height from the ground, but we do not analyse these data here.

We returned to our transect regularly (but not quite daily), to locate and measure new individuals and re-measure previously marked individuals. Not all individuals were located at the same time and so were not measured on the same days. When re-measuring individuals, we performed both behavioural tests and measures of the web described above, but we did not re-measure their length, as we assumed it was relatively invariant at the time scale we were working. Individuals’ web structures and behaviours were measured on average of 4.75 times (range 1-11, Fig. 1c). If we could not see an individual, we removed the flagging type and recorded it as gone. If an individual was found but had spun no orb that day, we did not measure its behaviour or its web structure, but also did not remove the flagging tape, allowing us to return and identify the indivdual. If an individual fled as we approached it, we measured its web structure and then left, returning later in the session to attempt to measure its behaviour. If we could not measure its behaviour that day, we recorded “NA” for both responsiveness to prey and reaction to a predation threat. If the spider fled from its web during the test for responsiveness to prey, we scored its responsiveness to prey as 180, and its reaction to a predation threat as 1. Our reasoning here is that a spider fleeing a potential prey item was both unresponsive to the opportunity and unwilling to face any potential predation risk. In five instances we re-tested such spiders once they had returned to the centre of the web, and in all cases they either fled again, or did not respond to the prey stimulus within 180 seconds, justifying our decision. We also tested whether these modelling decisions influenced our results, see “Robustness to modelling decisions”. In total we recorded 45 measures of length, 188 measures of responsiveness to a prey stimulus (146 of which were directly observed rather than assigned due to fleeing), 194 measures of reaction to a predation threat (146 of which were directly observed rather than assigned due to fleeing), 200 diameters, and 197 measures of web density and symmetry across 55 unique individuals (see Table 1 for means and variance of each trait).

**Table 1.**
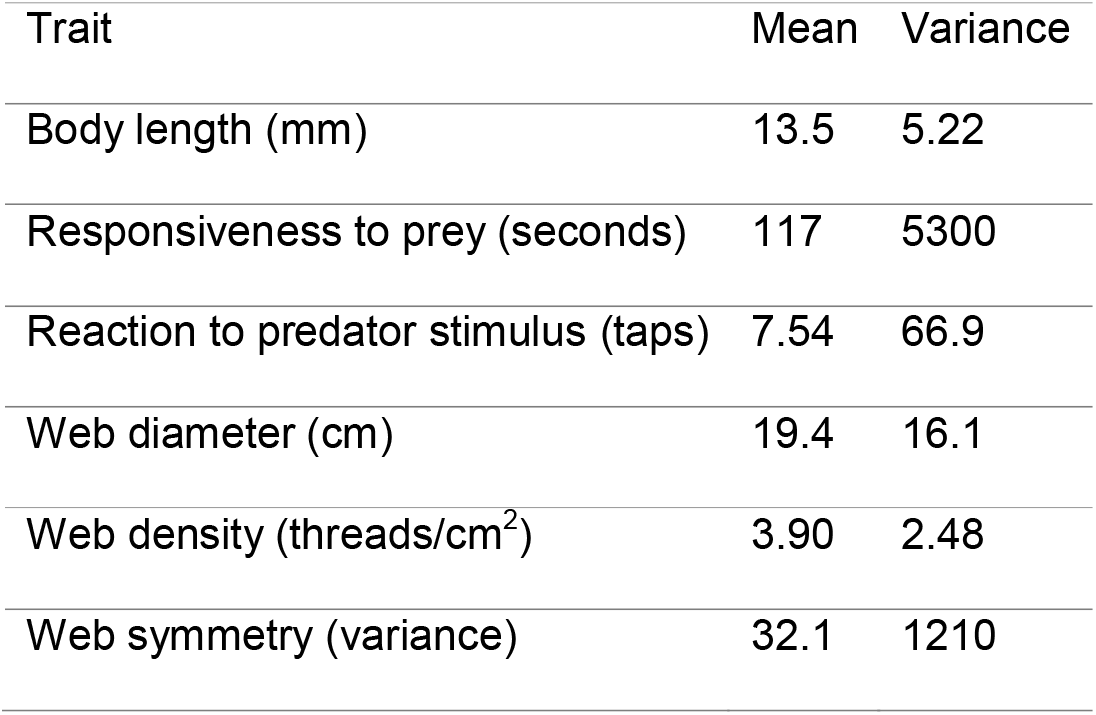
The means and variances of each of the six traits (to three significant figures). Note that web symmetry was log-transformed prior to analysis, while this logged variable, body length, web diameter and web evenness were mean centred and variance standardised prior to being entered into the analysis.

### Data analysis

We first estimated the phenotypic correlations between each trait pair (Pearson’s correlations; Fig. 2). To partition these correlations to the among-individual, among-date, and residual levels, we built a multivariate mixed model with each of the six traits as response variables. Responsiveness to prey and reaction to predation used a Poisson error distribution (log-link), while each web structure trait and body length used a Gaussian error distribution. Web symmetry was log-transformed, and then this transformed variable, web diameter, web density, and body length were mean centred and variance standardised (Schielzeth 2010). We fitted the random effect of individual identity and estimated the among-individual covariances among all six traits. We fitted the random effect of date and estimated the among-date covariances among all traits except for length, which was only measured on a single day per spider, and so we fixed the among-date variance to 0.0001. We estimated the residual covariances between all traits, and also fixed the residual variance for length to 0.0001, as it is only measured once per individual, following Houslay and Wilson(2017). We estimated unique intercepts for each response variable. Spiders might adjust their web structure or foraging behaviour over time as they gain information about their foraging patch (Nakata and Ushimaru 1999). Therefore, we fitted trial number (mean centred) as a fixed effect for each trait except length (which was only measured **once). The model was fitted in R (ver. 3.5.3; R Development Core Team 2016) with the** package “MCMCglmm” (Hadfield 2010). We used 550,000 iterations, a burn in of 50,000, and a thinning interval of 100. Priors were set to be flat and relatively uninformative, with 70% of the phenotypic variance for the logged values of each trait placed on the residual variance, 20% on the among-colony variance, and 10% on the among-date variance, following Brommer (2017). We calculated adjusted repeatabilities (after accounting for trial number) following Nakagawa and Schielzeth (2010).

**Figure 2.**
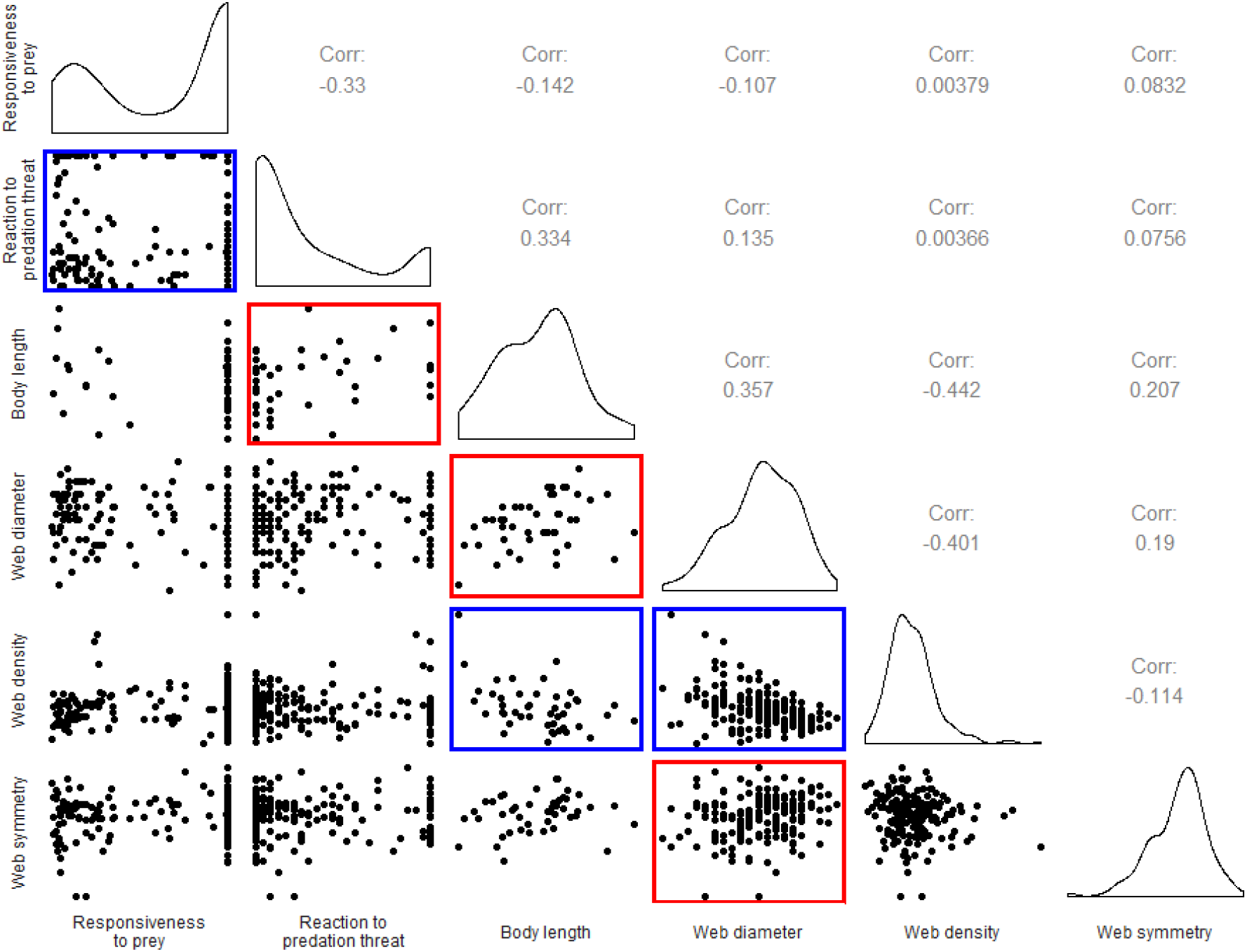
Phenotypic relationships between each of our six traits (web symmetry has been log-transformed). Pearson’s correlations are shown above the diagonal, pairwise plots below the diagonal, and histograms of each variable along the diagonal. Pairwise plots are bordered with red if the correlation was significant and positive, or bordered with blue if significant and negative.

### Robustness to modelling decisions

We repeated the analysis with all behavioural scores that were assigned when an individual fled rather than were directly observed set to “NA”. We also repeated this analysis with all behavioural scores that were ceiling values (180 for responsiveness to prey, 24 for reaction to a predation threat) set to “NA”. In each case our results did not change (full results of the original model are given in Table S1, with these auxiliary results shown in Tables S2&3). We therefore concluded that assigning behavioural scores to spiders that fled and giving unresponsive spiders the maximum score for each behaviour had not biased our results. We also repeated the original analysis without estimating any among-date covariances (as none were different from zero; see Table S1 & Fig. S1). This too did not change our results (Table S4), indicating that estimating the among-date covariances had not reduced our power and prevented us from detecting any other among-individual or residual covariances. As such we discuss the results as from the model where the among-date covariances were estimated.

## Results

### Repeatabilities

Web diameter was repeatable (r = 0.427, credible intervals [CIs] = 0.239 to 0.587), as was web density (r = 0.488, CIs = 0.286 to 0.646). The response to the predator stimulus was slightly repeatable (r = 0.045, CIs = < 0.001 to 0.159). Web symmetry was not repeatable (r < 0.001, CIs < 0.001 to 0.145), nor was response to the prey stimulus (r < 0.001, CIs < 0.001 to 0.012). Estimates for each of the among-individual, among-date, and residual variances for each trait are shown in Fig. 3. Individuals tended to react more quicklyto the predation threat in later trials, although this effect marginally overlapped with zero (fixed effect of trial, mode = −0.117, CIs = −0.252 to 0.011). Trial number did not influence any other trait.

**Figure 3.**
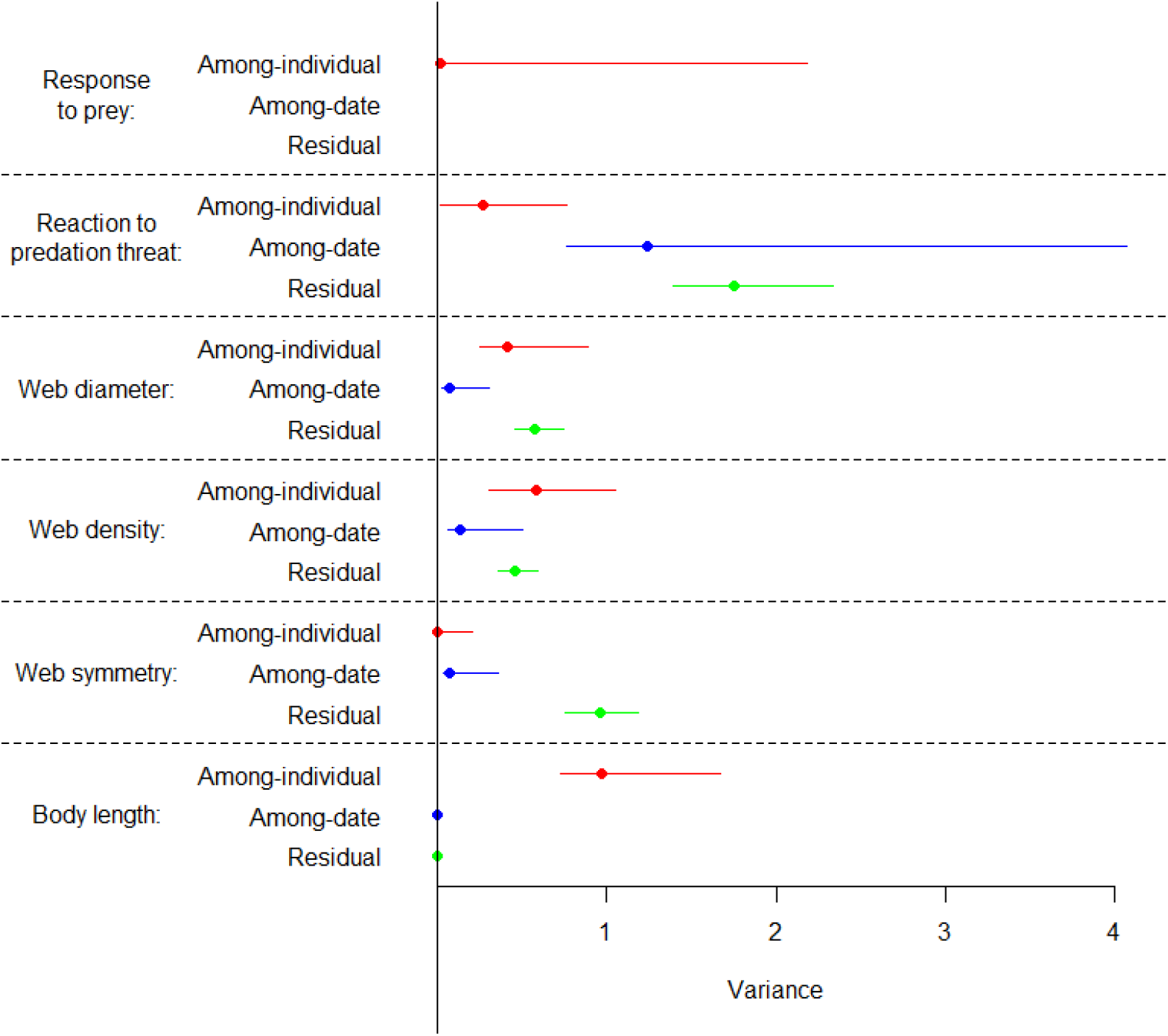
Variance in each trait partitioned to the among-individual (red), among-date (blue), or residual (green) levels. The among date variance for response to the prey stimulus was 87, while the residual variance was 44, and so are not shown on this plot. Both the among-date and residual variance for length were supressed to 0.0001.

### Covariances

Body length, web diameter and web density were associated into a syndrome at the among-individual level, such that longer spiders had wider and less dense webs (body length-web diameter correlation = 0.404, CIs = 0.130 to 0.708; body length-web density correlation = −0.608, CIs = −0.769 to −0.305, web diameter-web density correlation = −0.689, CIs = −0.865 to −0.309). Reaction to the predation threat tended to be associated with these traits as well, such spiders that reacted more slowly to the predation threat had longer bodies, and tended to have wider webs and less dense webs, but the latter two of these correlations overlapped with zero (reaction to predation-body length correlation = 0.620, CIs = 0.099 to 0.906; reaction to predation-web diameter correlation = 0.554, CIs = −0.104 to 0.998; reaction to predation-web density correlation = −0.744, CIs = −0.990 to 0.079). Plots of the estimated among-individual relationships are shown in Fig. 4, with estimates for the among-individual correlations shown in Fig. 5. Response to the prey stimulus and web symmetry were not associated with the other traits among individuals (Figs. 4 & 5).

**Figure 4.**
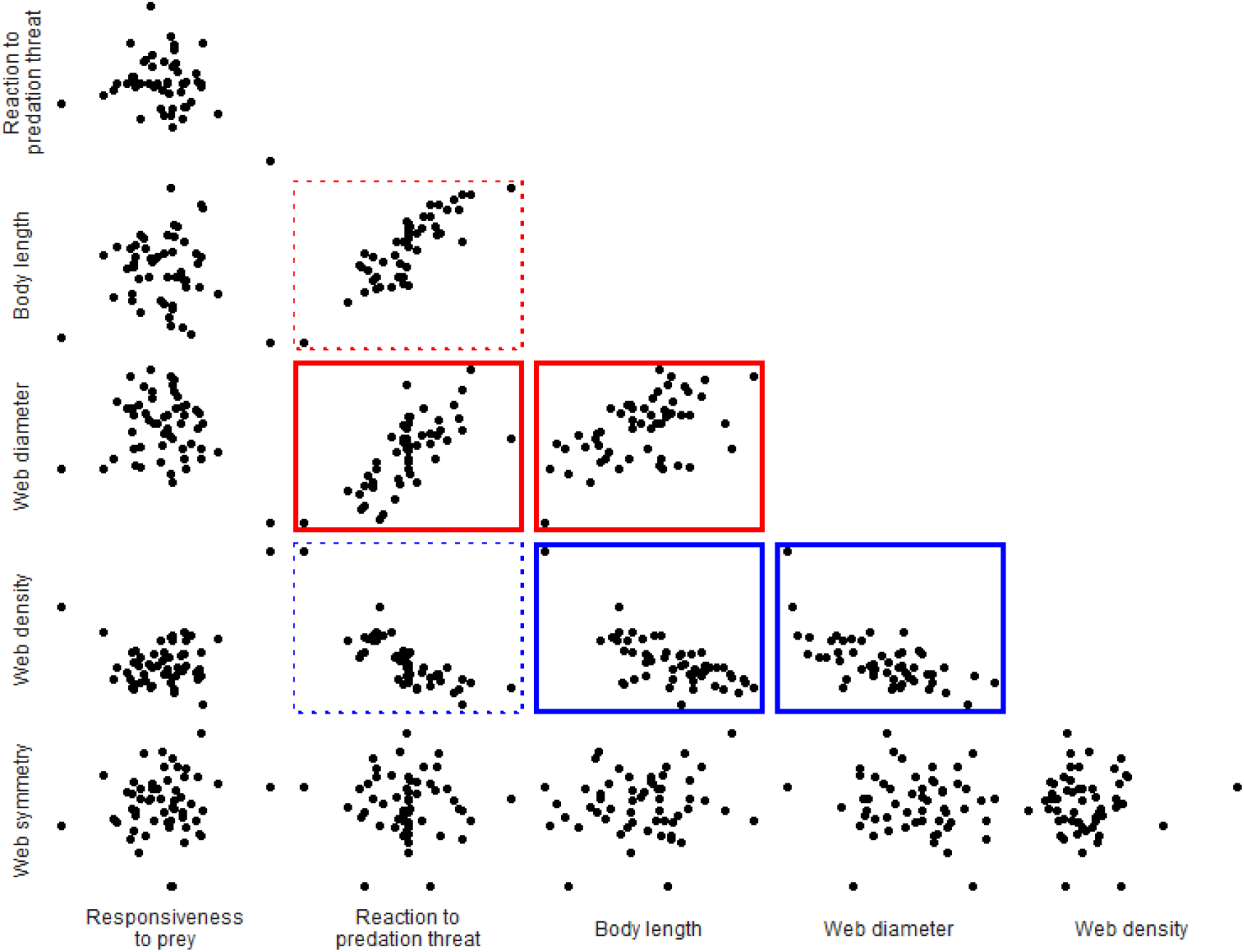
Pairwise plots of the estimated among-individual relationships between the six traits we studied, using best linear unbiased predictors extracted from the multivariate model. Pairwise plots are bordered with red if the correlation was significant and positive, or bordered with blue if significant and negative. The 95% credible intervals of the among-individual correlations between reaction to predation and both web diameter and web density overlapped zero, and so the borders are plotted with dashed lines.

**Figure 5.**
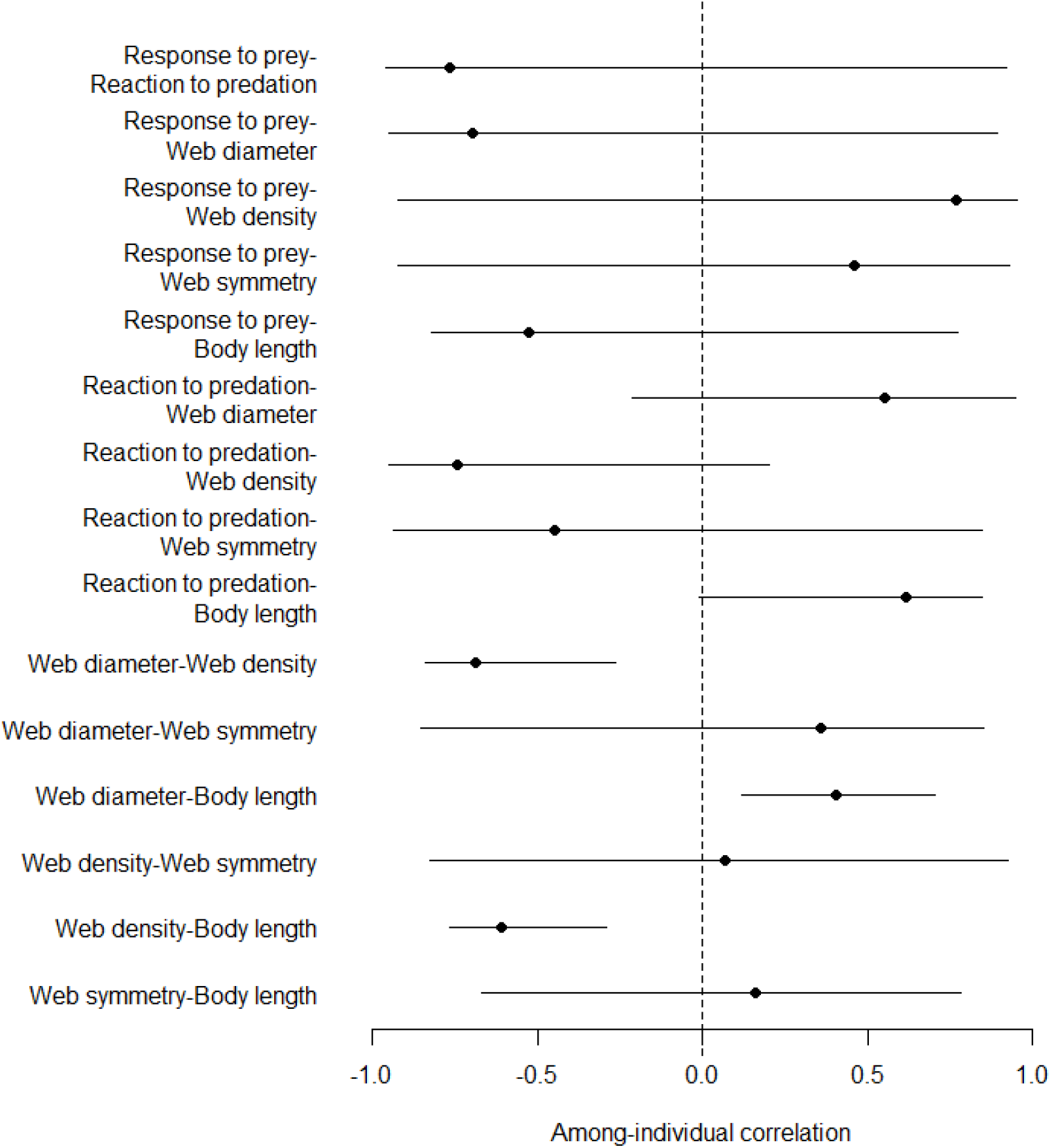
Estimated modes and 95% credible intervals of the among-individual correlations.

At the residual level there were associations between web diameter, density and symmetry, such that when a spider built a relatively wider web it also built a relatively less symmetrical and less dense web (Fig. 6; web diameter-web density correlation = −0.303, CIs = −0.438 to −0.115; web diameter-web symmetry correlation = 0.245, CIs = 0.073 to 0.379; web density-web symmetry correlation = −0.277, CIs = −0.442 to −0.139). No other residual covariances were different from zero (Fig. 6). There was variation among days for responsiveness to prey, reaction to predation, and somewhat so for web density (Fig. 3), but no among-date covariances were different from zero (Fig. S1).

**Figure 6.**
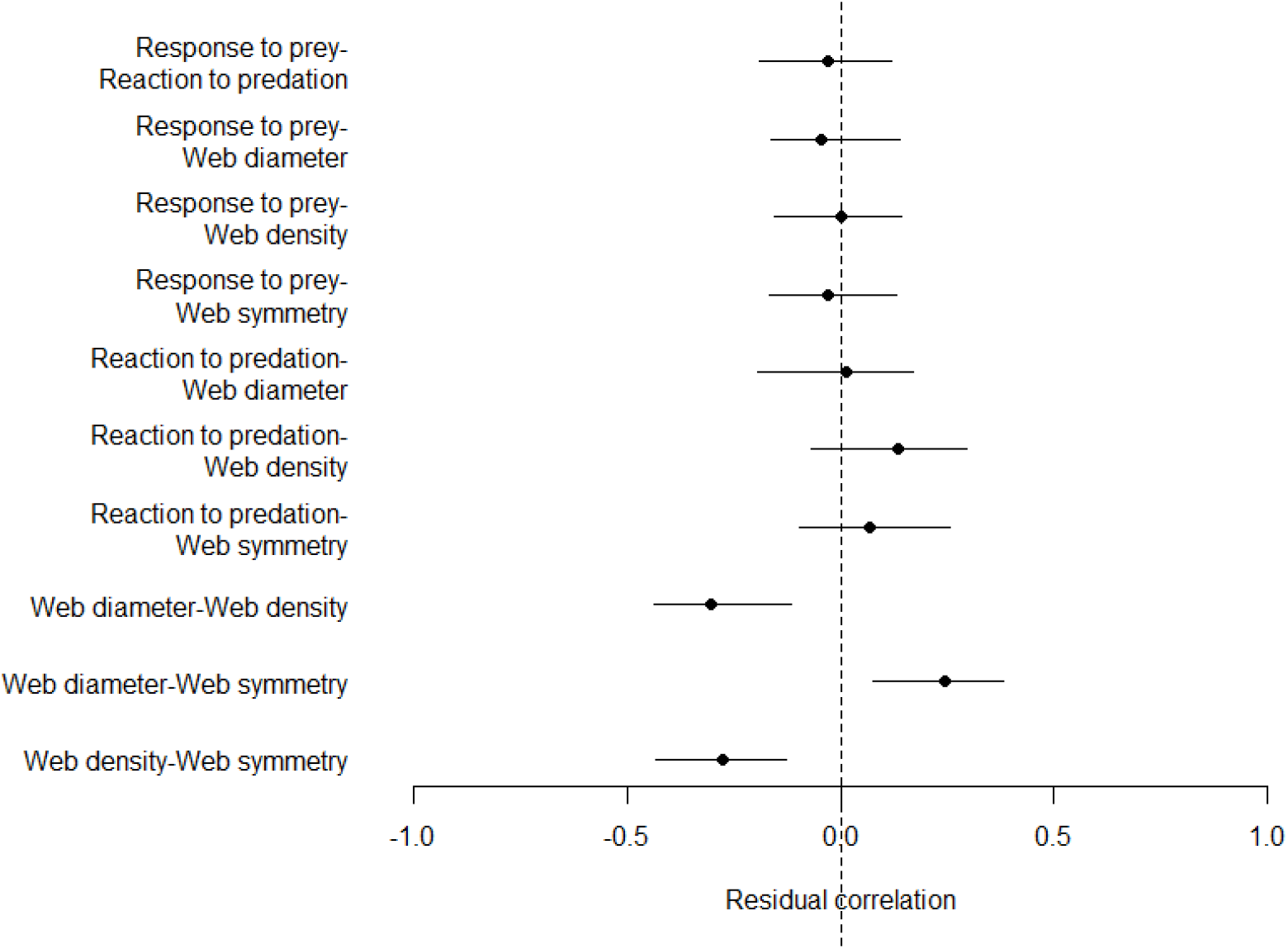
Estimated modes and 95% credible intervals of the residual correlations. Note that as the residual variance of body length was suppressed to 0.0001 it cannot covary at the residual level with the other traits, and so is not plotted here.

## Discussion

Individuals may show consistent differences in aspects of their behaviour, but whether their extended phenotypes are repeatable and covary with other behavioural traits is not well studied. We found that individual *Micrathena vigorsii* have consistent differences in their web diameters and web densities. These two web traits are correlated among-individuals into a syndrome with body length, and possibly reaction to a predation stimulus, indicating they might represent a single axis of variation. Responsiveness to a prey stimulus and web symmetry were not repeatable among-individuals and so were not associated with any traits among individuals.

Discovering a syndrome between web structure traits, morphology, and possibly behaviour suggests they may function together as a single integrated whole (e.g. they are an “evolutionary character”; Araya-Ajoy and Dingemanse 2014). This then raises questions as to the syndrome’s functional role, and the processes maintaining variation in this syndrome among-individuals (Sih, Bell, Johnson, et al. 2004; Sih, Bell, and Johnson 2004; Araya-Ajoy and Dingemanse 2014). The relationship between body length and web diameter suggests a simple biomechanical relationship between the size of spider (and perhaps the size of the steps it takes) and the resulting web it spins. This may naturally cause a wider web to be less dense, as a similar amount of silk is then spread over a larger area. However, if the relationship between web diameter and density was purely due to silk limitation, we would expect to see this trade-off at each level (among-date and residual) as well as at the among-individual level. Yet, we did not. This hypothesis also does not explain why reaction to a predation threat might be associated into the syndrome. Instead, variation in this size and stubbornness syndrome may represent an active strategy by larger individuals to catch larger and more nutritious, but possibly more dangerous, prey.

The density of a web may influence the type or size of prey it captures (Uetz et al. 1978; Chacon and Eberhard 1980; Blackledge and Zevenbergen 2006). Meanwhile, larger and more stubborn spiders might be willing to tackle larger and more dangerous prey, which smaller and more docile spiders would not risk (Mukherjee and Heithaus 2013; although juveniles in some species may be forced to target sub-optimal, and possibly more dangerous, prey; Elbroch et al. 2017). If body size and web size do relate to the type of prey captured, variation in the syndrome we have detected here might instead represent variation in individual foraging strategies (Bolnick et al. 2002; Ingram et al. 2018). In which case, we might expect the types and sizes of the prey caught in individual *M. vigorsii* webs to show consistent among-individual variation; a prediction that requires testing.

To predict how the syndrome might evolve, we need to know the degree to which it is heritable, and whether it is under selection. Around 50% of among-individual variation in behaviours may be based on additive genetic variance (Dochtermann et al. 2015), and genetic correlations tend to have the same sign as underlying genetic correlations (Dochtermann 2011) and so we could expect some of the (co)variation among-individuals in web structure and body length to have a genetic basis. In terms of selection, for now we do not have any estimates available for *M. vigorsii*. The different strategies may have equal pay-offs, which would maintain variation even if the syndrome had a genetic basis (Mangel and Stamps 2001; Stamps 2007). Alternatively, spiders may change their position in the syndrome as they age, spinning wider and less dense webs (and possibly reacting less to predation) as they grow. For example, in the orb-web spider *Zygiella x-notata*, webs have a shorter total thread length, become less regular, and have more “anomalies” as individuals age (Anotaux et al. 2012; Anotaux et al. 2014), and these changes increase prey handling time (Anotaux et al. 2014). We could not detect this trend in our study, as it took place at a time scale (one month, although each individual was tracked for considerably less time than this) where we might not expect to see much measurable growth. Determining whether *M. vigorsii* move through the size and stubbornness syndrome as they grow would require a much longer study, and one probably performed in captivity to track individuals over their entire lifetimes.

Some of the consistent variation in web structure we observed could be due to by variation in microhabitat, as in our study individuals were only ever assayed in one environment. Some variation in the amount of space a spider had to spin a web, the available structural supports, or some microclimatic factor, could cause consistent differences in web structure among-individuals (Zevenbergen et al. 2008; Blamires 2010; Nakata 2012). However, DiRienzo and Montiglio (Dirienzo and Montiglio 2016; Montiglio and DiRienzo 2016) also found consistent differences among-individuals in spider web structure, but in a laboratory setting. Such a setting should hypothetically control for variation in microhabitat, and so microhabitat variation could not be an explanation for consistent differences in individual web structure in their study; raising the possibility that microhabitat use may not fully explain our results either. Furthermore, a spider may select the microhabitat that allows the spinning of a web of a certain structure (see also “niche construction” Odling-Smee et al. 2003; Saltz and Nuzhdin 2014). Therefore, while web structure and microhabitat could covary, this could still depend on the spider’s decision making, and so would still be classed as a trait of the spider, not as one driven by the environment.

Our results are generally in agreement with previous work. DiRienzo and Montiglio (2016) found consistent differences among-individuals in web structure, and that black widow spiders (*Latrodectus hesperus*) with longer femur–patellas build webs with more gumfooted lines. We also found consistent differences among-individuals in web structure, and a positive relationship between body size and web size. We therefore tentatively suggest that these two elements could be general features of intra-population variation in spider webs. More studies in other taxa with different extended phenotypes are required to determine whether, within a population, larger individuals usually build bigger nests or larger dams, and so on. DiRienzo and Montilgio (2016) also found that a higher number of gumfooted lines is associated with increased foraging aggression. However, Montiglio and DiRienzo (2016), also in *L. hesperus*, found a higher number of both gumfooted and structural lines is associated with *decreased* foraging aggression, while boldness was not associated with any web characteristics. Given that the relationship between behaviour and web structure we detected overlapped with zero, and that responsiveness to a prey stimulus was not associated with web structure, in aggregate it seems there is yet no clear pattern in how web structure and behaviours are associated among-individuals.

The residual covariances between web diameter, web density, and web symmetry indicates that when a spider builds a relatively larger web (relative to its own typical web diameter) it also builds a relatively less symmetrical and less dense web. A spider may struggle to apply its usual web spinning strategy at a greater spatial scale without making mistakes, and so created a less symmetrical and less dense web when it tries to make a larger web than usual. Alternatively, our measure of web symmetry could reflect the amount of contact with prey the web experienced that day (webs are typically spun in the morning, and all web measurements were taken in the afternoon). *M. vigorsii* do not leave prey remains in their web (DN Fisher, pers obs.), but contact with prey, whether leading to a successful capture or not, leads to the removal of spirals and radii. The removal of these threads reduces density, as well as change the number of spirals along one axis but not others, giving the impression of a less symmetrical web. A greater rate of contact with prey would be expected with larger webs as they cover a greater area, but we would perhaps expect this covariance to be present at the among-individual and among-date levels as well as at the residual level. We therefore cannot identify what process might be driving the residual covariances between web diameter, density, and symmetry at this time.

We note here that correlated measurement error can give residual covariances between traits. We think this is unlikely to have occurred here, as web diameter and the other measures of web structure were measured using different tools, and web density and web symmetry were calculated once we had returned from the field. This reduces the chance that we could make mistakes that simultaneously influenced all measurements. Further, previous work has suggested that spiders spin webs suited for defence rather than prey capture, and respond less to prey, following prey capture (Venner et al. 2000; Dirienzo and Montiglio 2016 Jul). This would appear as a residual correlation between web diameter, web density and responsiveness to prey in our analysis, but responsiveness to prey did not covary with these traits at the residual (or any) level. This suggests *M. vigorsii* does not simply reduce the investment in its web and its responsiveness to prey following a day where it captured prey, though this would need to be confirmed experimentally.

There was a phenotypic correlation between responsiveness to prey and reaction to predation (Fig. 2) that was not different from zero as an among-individual correlation (or indeed at any other level; Figs. 4–6 & S1). This acts as a cautionary tale against assuming any phenotypic correlations represent among-individual correlations (the “individual gambit”; Brommer 2013). In general, we should be careful to identify at what level (within-individual, among-individual, among-day) any hypothesised relationships between traits should exist, and then construct statistical models that specifically estimate these terms, allowing us to directly evaluate our hypotheses (Dingemanse and Dochtermann 2013; e.g. Moiron et al. 2016).

## Conclusions

We found that *M. vigorsii* show consistent among-individual differences in aspects of web structure, and that there is a syndrome between web structure and morphology such that larger individuals spin wider and less dense webs. These may be general features of intra-population (co)variation in morphology and extended phenotypes and could represent among-individual variation in foraging strategies or aging-dependent changes in various aspects of the phenotype. Reaction to a predator stimulus was very slightly repeatable and this trait may be integrated into the among-individual syndrome with morphology and web structure, but uncertainty was high. These results highlight how extended phenotypes can be integrated into general suites of trait variation, and so selection likely acts upon these traits in concert. We also found that when a spider builds a relatively wider web the web is also relatively less dense and less symmetrical, suggesting spiders building relatively larger webs make more mistakes than when building a web closer to the average size for that individual. Responsiveness to a prey stimulus and web symmetry were not consistently different among-individuals, and so are completely plastic or environmentally determined traits that are not based on more stable individual characteristics. Extended phenotypes like the web traits evaluated here represent a suite of biological traits that have perhaps been under-studied in the past literature on individual variation, yet these traits represent important biological variation that can play key roles in organisms’ lives and in ecosystems more broadly. Here we have demonstrated that, in a population of wild organisms, aspects of an extended phenotype are integrated into a syndrome with morphological, and possibly behavioural, traits. Therefore, our expectations for selection on and evolution of extended phenotypes should consider how they integrate with other phenotypes under natural conditions.

## Author contributions

DNF and JNP designed the study. JY acquired the permits. DNF collected and analysed the data and drafted the manuscript. All authors contributed to revisions of the manuscript and approved the final version.

## Acknowledgements

We thank J. B. Barnett, H. M. Anderson, B. L. McEwen, J. L. L. Lichtenstein, and R. Costa-Pereira for being excellent colleagues in the field. We also thank T. D. Swanson and the staff of the Andes and Amazon Field School at Iyarina for making our stay as comfortable and enjoyable as possible. Funding was provided by a Canada 150 Research Chair award to JNP. We have no conflicts of interest.

## Data accessibility

All data and R code used in the analysis will be made available upon publication.

