## Supplementary materials – Fig S1 & Tables S1-4 for "Orb-weaving spiders show a correlated syndrome of morphology and web structure in the wild"

Supplementary figure

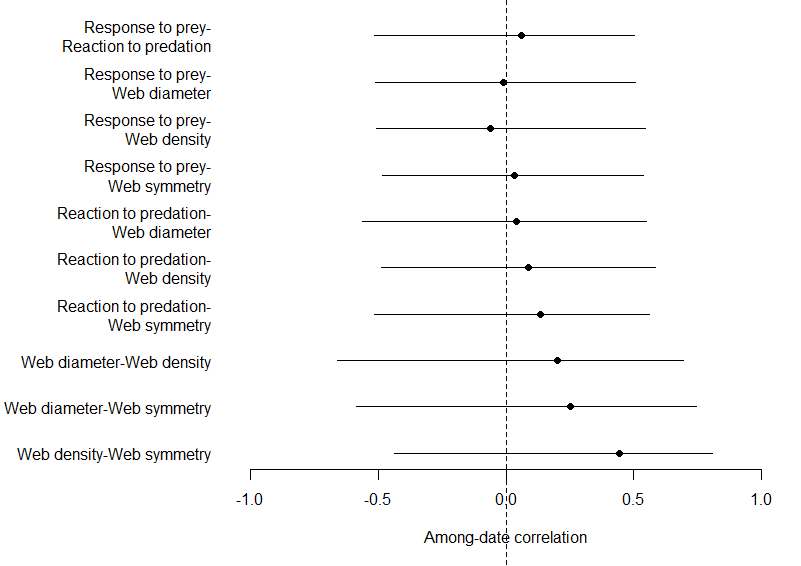

**Figure S1**. Estimated modes and 95% credible intervals of the among-date correlations from the original model in the main text (see also Table S1). Note that as the among- date variance of body length was suppressed to 0.0001 it cannot covary at the among-date level with the other traits, and so is not plotted here.

Supplementary tables

**Table S1**. Model results for original multivariate model in main text

| Term | Trait(s) | Posterior mean | Lower 95% CI | Upper 95% CI |
| --- | --- | --- | --- | --- |
| Intercept | Response to prey | 2.650 | -1.227 | 6.591 |
| Intercept | Reaction to predation threat | 1.305 | 0.603 | 2.040 |
| Intercept | Web diameter | -0.038 | -0.331 | 0.237 |
| Intercept | Web density | 0.065 | -0.279 | 0.372 |
| Intercept | Web symmetry | 0.004 | -0.246 | 0.242 |
| Intercept | Body length | 0.021 | -0.269 | 0.293 |
| Trial | Response to prey | -0.008 | -0.587 | 0.549 |
| Trial | Reaction to predation threat | -0.110 | -0.240 | 0.019 |
| Trial | Web diameter | 0.006 | -0.063 | 0.065 |
| Trial | Web density | -0.020 | -0.084 | 0.049 |
| Trial | Web symmetry | 0.002 | -0.063 | 0.070 |
| Trial | Body length | 0.00000 | -0.0001 | 0.0001 |
| Among-individual variance | Response to prey | 0.423 | 0.00000 | 1.632 |
| Among-individual covariance | Response to prey – Reaction to predation threat | -0.062 | -0.532 | 0.335 |
| Among-individual covariance | Response to prey – Web diameter | -0.081 | -0.701 | 0.442 |
| Among-individual covariance | Response to prey – Web density | 0.074 | -0.592 | 0.757 |
| Among-individual covariance | Response to prey – Web symmetry | 0.003 | -0.165 | 0.182 |
| Among-individual covariance | Response to prey – Body length | -0.092 | -0.880 | 0.571 |
| Among-individual variance | Reaction to predation threat | 0.298 | 0.00000 | 0.667 |
| Among-individual covariance | Reaction to predation threat – Web diameter | 0.166 | -0.097 | 0.417 |
| Among-individual covariance | Reaction to predation threat – Web density | -0.198 | -0.501 | 0.087 |
| Among-individual covariance | Reaction to predation threat – Web symmetry | -0.024 | -0.163 | 0.077 |
| Among-individual covariance | Reaction to predation threat – Body length | 0.287 | -0.027 | 0.623 |
| Among-individual variance | Web diameter | 0.499 | 0.206 | 0.825 |
| Among-individual covariance | Web diameter – Web density | -0.334 | -0.614 | -0.083 |
| Among-individual covariance | Web diameter – Web symmetry | 0.012 | -0.116 | 0.152 |
| Among-individual covariance | Web diameter – Body length | 0.325 | 0.070 | 0.616 |
| Among-individual variance | Web density | 0.611 | 0.273 | 0.984 |
| Among-individual covariance | Web density – Web symmetry | 0.012 | -0.137 | 0.165 |
| Among-individual covariance | Web density – Body length | -0.458 | -0.778 | -0.184 |
| Among-individual variance | Web symmetry | 0.057 | 0 | 0.170 |
| Among-individual covariance | Web symmetry – Body length | 0.036 | -0.126 | 0.224 |
| Among-individual variance | Body length | 1.096 | 0.681 | 1.576 |
| Among-date variance | Response to prey | 105.804 | 34.277 | 198.024 |
| Among-date covariance | Response to prey – Reaction to predation threat | -0.087 | -8.506 | 9.102 |
| Among-date covariance | Response to prey – Web diameter | 0.001 | -2.239 | 2.047 |
| Among-date covariance | Response to prey – Web density | 0.105 | -2.726 | 3.238 |
| Among-date covariance | Response to prey – Web symmetry | 0.117 | -2.393 | 2.461 |
| Among-date covariance | Response to prey – Body length | 0.0001 | -0.017 | 0.018 |
| Among-date variance | Reaction to predation threat | 1.807 | 0.635 | 3.484 |
| Among-date covariance | Reaction to predation threat – Web diameter | -0.003 | -0.302 | 0.299 |
| Among-date covariance | Reaction to predation threat – Web density | 0.040 | -0.331 | 0.471 |
| Among-date covariance | Reaction to predation threat – Web symmetry | 0.017 | -0.283 | 0.356 |
| Among-date covariance | Reaction to predation threat – Body length | 0.00000 | -0.003 | 0.002 |
| Among-date variance | Web diameter | 0.109 | 0.016 | 0.243 |
| Among-date covariance | Web diameter – Web density | -0.001 | -0.138 | 0.124 |
| Among-date covariance | Web diameter – Web symmetry | 0.017 | -0.087 | 0.128 |
| Among-date covariance | Web diameter – Body length | -0.00000 | -0.001 | 0.001 |
| Among-date variance | Web density | 0.195 | 0.045 | 0.414 |
| Among-date covariance | Web density – Web symmetry | 0.047 | -0.089 | 0.189 |
| Among-date covariance | Web density – Body length | 0.00001 | -0.001 | 0.001 |
| Among-date variance | Web symmetry | 0.135 | 0.020 | 0.304 |
| Among-date covariance | Web symmetry – Body length | -0.00000 | -0.001 | 0.001 |
| Among-date variance | Body length | 0.00001 | 0.00001 | 0.00001 |
| Residual variance | Response to prey | 45.876 | 36.227 | 55.601 |
| Residual covariance | Response to prey – Reaction to predation threat | -0.336 | -1.751 | 1.083 |
| Residual covariance | Response to prey – Web diameter | -0.063 | -0.818 | 0.730 |
| Residual covariance | Response to prey – Web density | -0.033 | -0.706 | 0.679 |
| Residual covariance | Response to prey – Web symmetry | -0.128 | -1.085 | 0.927 |
| Residual covariance | Response to prey – Body length | 0.0001 | -0.008 | 0.009 |
| Residual variance | Reaction to predation threat | 1.818 | 1.387 | 2.320 |
| Residual covariance | Reaction to predation threat – Web diameter | -0.012 | -0.198 | 0.180 |
| Residual covariance | Reaction to predation threat – Web density | 0.106 | -0.059 | 0.288 |
| Residual covariance | Reaction to predation threat – Web symmetry | 0.108 | -0.130 | 0.337 |
| Residual covariance | Reaction to predation threat – Body length | 0.00003 | -0.003 | 0.003 |
| Residual variance | Web diameter | 0.580 | 0.438 | 0.721 |
| Residual covariance | Web diameter – Web density | -0.146 | -0.238 | -0.049 |
| Residual covariance | Web diameter – Web symmetry | 0.175 | 0.049 | 0.301 |
| Residual covariance | Web diameter – Body length | 0.00001 | -0.004 | 0.004 |
| Residual variance | Web density | 0.463 | 0.352 | 0.580 |
| Residual covariance | Web density – Web symmetry | -0.191 | -0.305 | -0.078 |
| Residual covariance | Web density – Body length | 0.00001 | -0.004 | 0.004 |
| Residual variance | Web symmetry | 0.954 | 0.753 | 1.175 |
| Residual covariance | Web symmetry – Body length | -0.00005 | -0.006 | 0.006 |
| Residual variance | Body length | 0.0001 | 0.0001 | 0.0001 |

**Table S2**. Model results for model where occasions where spiders fled during the prey test, and so were assigned “NAs” for responsiveness to prey and Reaction to predation threat rather than 180 and 24 respectively.

| Term | Trait(s) | Posterior mean | Lower 95% CI | Upper 95% CI |
| --- | --- | --- | --- | --- |
| Intercept | Response to prey | 2.182 | -1.977 | 6.217 |
| Intercept | Reaction to predation threat | 1.766 | 1.062 | 2.520 |
| Intercept | Web diameter | -0.036 | -0.330 | 0.246 |
| Intercept | Web density | 0.057 | -0.270 | 0.385 |
| Intercept | Web symmetry | 0.003 | -0.242 | 0.252 |
| Intercept | Body length | 0.045 | -0.231 | 0.327 |
| Trial | Response to prey | 0.025 | -0.659 | 0.818 |
| Trial | Reaction to predation threat | -0.121 | -0.260 | 0.014 |
| Trial | Web diameter | 0.007 | -0.057 | 0.067 |
| Trial | Web density | -0.022 | -0.090 | 0.042 |
| Trial | Web symmetry | -0.001 | -0.065 | 0.070 |
| Trial | Body length | 0.00000 | -0.0001 | 0.0001 |
| Among-individual variance | Response to prey | 0.671 | 0.00000 | 2.614 |
| Among-individual covariance | Response to prey – Reaction to predation threat | -0.023 | -0.378 | 0.331 |
| Among-individual covariance | Response to prey – Web diameter | -0.071 | -0.933 | 0.665 |
| Among-individual covariance | Response to prey – Web density | 0.066 | -0.769 | 1.121 |
| Among-individual covariance | Response to prey – Web symmetry | 0.001 | -0.193 | 0.218 |
| Among-individual covariance | Response to prey – Body length | -0.066 | -1.038 | 0.855 |
| Among-individual variance | Reaction to predation threat | 0.100 | 0 | 0.318 |
| Among-individual covariance | Reaction to predation threat – Web diameter | 0.108 | -0.066 | 0.346 |
| Among-individual covariance | Reaction to predation threat – Web density | -0.105 | -0.373 | 0.102 |
| Among-individual covariance | Reaction to predation threat – Web symmetry | -0.003 | -0.082 | 0.064 |
| Among-individual covariance | Reaction to predation threat – Body length | 0.138 | -0.088 | 0.430 |
| Among-individual variance | Web diameter | 0.520 | 0.232 | 0.854 |
| Among-individual covariance | Web diameter – Web density | -0.356 | -0.638 | -0.095 |
| Among-individual covariance | Web diameter – Web symmetry | 0.011 | -0.121 | 0.147 |
| Among-individual covariance | Web diameter – Body length | 0.356 | 0.095 | 0.679 |
| Among-individual variance | Web density | 0.635 | 0.292 | 1.030 |
| Among-individual covariance | Web density – Web symmetry | 0.011 | -0.130 | 0.174 |
| Among-individual covariance | Web density – Body length | -0.470 | -0.795 | -0.193 |
| Among-individual variance | Web symmetry | 0.047 | 0 | 0.156 |
| Among-individual covariance | Web symmetry – Body length | 0.042 | -0.108 | 0.231 |
| Among-individual variance | Body length | 1.110 | 0.704 | 1.578 |
| Among-date variance | Response to prey | 112.926 | 40.013 | 210.208 |
| Among-date covariance | Response to prey – Reaction to predation threat | 0.021 | -9.823 | 8.709 |
| Among-date covariance | Response to prey – Web diameter | 0.028 | -2.144 | 2.357 |
| Among-date covariance | Response to prey – Web density | 0.057 | -3.032 | 3.276 |
| Among-date covariance | Response to prey – Web symmetry | 0.075 | -2.390 | 2.772 |
| Among-date covariance | Response to prey – Body length | -0.00005 | -0.019 | 0.018 |
| Among-date variance | Reaction to predation threat | 1.862 | 0.612 | 3.626 |
| Among-date covariance | Reaction to predation threat – Web diameter | -0.001 | -0.333 | 0.281 |
| Among-date covariance | Reaction to predation threat – Web density | 0.048 | -0.383 | 0.443 |
| Among-date covariance | Reaction to predation threat – Web symmetry | 0.044 | -0.260 | 0.447 |
| Among-date covariance | Reaction to predation threat – Body length | 0.00000 | -0.002 | 0.003 |
| Among-date variance | Web diameter | 0.104 | 0.020 | 0.231 |
| Among-date covariance | Web diameter – Web density | 0.004 | -0.119 | 0.116 |
| Among-date covariance | Web diameter – Web symmetry | 0.017 | -0.090 | 0.119 |
| Among-date covariance | Web diameter – Body length | 0.00000 | -0.001 | 0.001 |
| Among-date variance | Web density | 0.186 | 0.043 | 0.385 |
| Among-date covariance | Web density – Web symmetry | 0.048 | -0.071 | 0.195 |
| Among-date covariance | Web density – Body length | 0.00000 | -0.001 | 0.001 |
| Among-date variance | Web symmetry | 0.138 | 0.027 | 0.298 |
| Among-date covariance | Web symmetry – Body length | -0.00000 | -0.001 | 0.001 |
| Among-date variance | Body length | 0.00001 | 0.00001 | 0.00001 |
| Residual variance | Response to prey | 61.460 | 46.594 | 78.720 |
| Residual covariance | Response to prey – Reaction to predation threat | 0.011 | -1.902 | 1.822 |
| Residual covariance | Response to prey – Web diameter | -0.067 | -1.173 | 1.004 |
| Residual covariance | Response to prey – Web density | 0.095 | -0.813 | 1.002 |
| Residual covariance | Response to prey – Web symmetry | -0.076 | -1.360 | 1.306 |
| Residual covariance | Response to prey – Body length | 0.0001 | -0.011 | 0.011 |
| Residual variance | Reaction to predation threat | 1.792 | 1.328 | 2.250 |
| Residual covariance | Reaction to predation threat – Web diameter | 0.012 | -0.199 | 0.212 |
| Residual covariance | Reaction to predation threat – Web density | 0.066 | -0.118 | 0.243 |
| Residual covariance | Reaction to predation threat – Web symmetry | 0.030 | -0.212 | 0.292 |
| Residual covariance | Reaction to predation threat – Body length | 0.00002 | -0.003 | 0.003 |
| Residual variance | Web diameter | 0.575 | 0.432 | 0.715 |
| Residual covariance | Web diameter – Web density | -0.139 | -0.238 | -0.051 |
| Residual covariance | Web diameter – Web symmetry | 0.170 | 0.043 | 0.299 |
| Residual covariance | Web diameter – Body length | -0.0001 | -0.004 | 0.004 |
| Residual variance | Web density | 0.455 | 0.343 | 0.565 |
| Residual covariance | Web density – Web symmetry | -0.184 | -0.298 | -0.072 |
| Residual covariance | Web density – Body length | 0.0001 | -0.004 | 0.004 |
| Residual variance | Web symmetry | 0.958 | 0.761 | 1.174 |
| Residual covariance | Web symmetry – Body length | 0.00001 | -0.006 | 0.006 |
| Residual variance | Body length | 0.0001 | 0.0001 | 0.0001 |

**Table S3**. Model results for model where occasions where spiders recorded maximum scores for responsiveness to prey and Reaction to predation threat (180 and 24 respectively) were assigned “NAs”.

| Term | Trait(s) | Posterior mean | Lower 95% CI | Upper 95% CI |
| --- | --- | --- | --- | --- |
| Intercept | Response to prey | 2.094 | -1.906 | 6.492 |
| Intercept | Reaction to predation threat | 1.020 | 0.286 | 1.715 |
| Intercept | Web diameter | -0.038 | -0.339 | 0.236 |
| Intercept | Web density | 0.074 | -0.272 | 0.387 |
| Intercept | Web symmetry | -0.005 | -0.259 | 0.233 |
| Intercept | Body length | -0.020 | -0.300 | 0.246 |
| Trial | Response to prey | 0.022 | -1.350 | 1.300 |
| Trial | Reaction to predation threat | -0.103 | -0.233 | 0.035 |
| Trial | Web diameter | 0.007 | -0.056 | 0.072 |
| Trial | Web density | -0.023 | -0.091 | 0.042 |
| Trial | Web symmetry | -0.0005 | -0.067 | 0.067 |
| Trial | Body length | -0.00000 | -0.0001 | 0.0001 |
| Among-individual variance | Response to prey | 2.049 | 0.00000 | 7.953 |
| Among-individual covariance | Response to prey – Reaction to predation threat | -0.053 | -0.716 | 0.458 |
| Among-individual covariance | Response to prey – Web diameter | -0.147 | -1.458 | 1.101 |
| Among-individual covariance | Response to prey – Web density | 0.168 | -1.316 | 1.750 |
| Among-individual covariance | Response to prey – Web symmetry | -0.011 | -0.419 | 0.347 |
| Among-individual covariance | Response to prey – Body length | -0.228 | -1.865 | 1.356 |
| Among-individual variance | Reaction to predation threat | 0.099 | 0 | 0.318 |
| Among-individual covariance | Reaction to predation threat – Web diameter | 0.056 | -0.125 | 0.273 |
| Among-individual covariance | Reaction to predation threat – Web density | -0.081 | -0.337 | 0.127 |
| Among-individual covariance | Reaction to predation threat – Web symmetry | 0.003 | -0.070 | 0.076 |
| Among-individual covariance | Reaction to predation threat – Body length | 0.145 | -0.075 | 0.440 |
| Among-individual variance | Web diameter | 0.506 | 0.219 | 0.841 |
| Among-individual covariance | Web diameter – Web density | -0.332 | -0.610 | -0.078 |
| Among-individual covariance | Web diameter – Web symmetry | 0.011 | -0.108 | 0.151 |
| Among-individual covariance | Web diameter – Body length | 0.274 | 0.029 | 0.539 |
| Among-individual variance | Web density | 0.605 | 0.279 | 0.991 |
| Among-individual covariance | Web density – Web symmetry | 0.009 | -0.128 | 0.163 |
| Among-individual covariance | Web density – Body length | -0.406 | -0.698 | -0.137 |
| Among-individual variance | Web symmetry | 0.047 | 0 | 0.151 |
| Among-individual covariance | Web symmetry – Body length | 0.059 | -0.084 | 0.237 |
| Among-individual variance | Body length | 1.008 | 0.634 | 1.438 |
| Among-date variance | Response to prey | 133.183 | 44.491 | 260.446 |
| Among-date covariance | Response to prey – Reaction to predation threat | -0.181 | -10.056 | 9.955 |
| Among-date covariance | Response to prey – Web diameter | 0.027 | -2.572 | 2.760 |
| Among-date covariance | Response to prey – Web density | -0.062 | -3.909 | 3.378 |
| Among-date covariance | Response to prey – Web symmetry | 0.066 | -3.029 | 2.973 |
| Among-date covariance | Response to prey – Body length | 0.00003 | -0.021 | 0.021 |
| Among-date variance | Reaction to predation threat | 1.824 | 0.595 | 3.514 |
| Among-date covariance | Reaction to predation threat – Web diameter | -0.029 | -0.333 | 0.338 |
| Among-date covariance | Reaction to predation threat – Web density | 0.059 | -0.327 | 0.498 |
| Among-date covariance | Reaction to predation threat – Web symmetry | 0.014 | -0.309 | 0.377 |
| Among-date covariance | Reaction to predation threat – Body length | 0.00001 | -0.002 | 0.003 |
| Among-date variance | Web diameter | 0.114 | 0.020 | 0.262 |
| Among-date covariance | Web diameter – Web density | -0.004 | -0.141 | 0.127 |
| Among-date covariance | Web diameter – Web symmetry | 0.018 | -0.084 | 0.129 |
| Among-date covariance | Web diameter – Body length | 0.00001 | -0.001 | 0.001 |
| Among-date variance | Web density | 0.198 | 0.050 | 0.425 |
| Among-date covariance | Web density – Web symmetry | 0.050 | -0.079 | 0.191 |
| Among-date covariance | Web density – Body length | 0.00001 | -0.001 | 0.001 |
| Among-date variance | Web symmetry | 0.135 | 0.021 | 0.301 |
| Among-date covariance | Web symmetry – Body length | -0.00000 | -0.001 | 0.001 |
| Among-date variance | Body length | 0.00001 | 0.00001 | 0.00001 |
| Residual variance | Response to prey | 103.825 | 74.023 | 139.844 |
| Residual covariance | Response to prey – Reaction to predation threat | 0.003 | -2.834 | 2.769 |
| Residual covariance | Response to prey – Web diameter | -0.177 | -1.858 | 1.576 |
| Residual covariance | Response to prey – Web density | 0.042 | -1.383 | 1.553 |
| Residual covariance | Response to prey – Web symmetry | 0.067 | -1.986 | 2.176 |
| Residual covariance | Response to prey – Body length | -0.0005 | -0.019 | 0.019 |
| Residual variance | Reaction to predation threat | 1.792 | 1.317 | 2.271 |
| Residual covariance | Reaction to predation threat – Web diameter | 0.048 | -0.147 | 0.244 |
| Residual covariance | Reaction to predation threat – Web density | 0.060 | -0.111 | 0.233 |
| Residual covariance | Reaction to predation threat – Web symmetry | -0.002 | -0.230 | 0.243 |
| Residual covariance | Reaction to predation threat – Body length | -0.00001 | -0.003 | 0.003 |
| Residual variance | Web diameter | 0.571 | 0.436 | 0.715 |
| Residual covariance | Web diameter – Web density | -0.140 | -0.232 | -0.045 |
| Residual covariance | Web diameter – Web symmetry | 0.170 | 0.046 | 0.300 |
| Residual covariance | Web diameter – Body length | -0.00000 | -0.004 | 0.004 |
| Residual variance | Web density | 0.457 | 0.350 | 0.573 |
| Residual covariance | Web density – Web symmetry | -0.185 | -0.288 | -0.069 |
| Residual covariance | Web density – Body length | 0.0001 | -0.004 | 0.004 |
| Residual variance | Web symmetry | 0.959 | 0.758 | 1.177 |
| Residual covariance | Web symmetry – Body length | -0.0003 | -0.006 | 0.006 |
| Residual variance | Body length | 0.0001 | 0.0001 | 0.0001 |

**Table S4**. Model results from the model where all among-date covariances were assumed to be zero and so not estimated.

| Term | Trait(s) | Posterior mean | Lower 95% CI | Upper 95% CI |
| --- | --- | --- | --- | --- |
| Intercept | Response to prey | 2.917 | -0.597 | 6.712 |
| Intercept | Reaction to predation threat | 1.303 | 0.729 | 1.916 |
| Intercept | Web diameter | -0.044 | -0.303 | 0.212 |
| Intercept | Web density | 0.064 | -0.219 | 0.358 |
| Intercept | Web symmetry | 0.001 | -0.218 | 0.206 |
| Intercept | Body length | 0.056 | -0.243 | 0.333 |
| Trial | Response to prey | -0.014 | -0.606 | 0.510 |
| Trial | Reaction to predation threat | -0.111 | -0.231 | 0.016 |
| Trial | Web diameter | 0.004 | -0.056 | 0.067 |
| Trial | Web density | -0.022 | -0.085 | 0.043 |
| Trial | Web symmetry | 0.002 | -0.062 | 0.066 |
| Trial | Body length | 0.00000 | -0.0001 | 0.0001 |
| Among-individual variance | Response to prey | 0.406 | 0.00000 | 1.561 |
| Among-individual covariance | Response to prey – Reaction to predation threat | -0.060 | -0.531 | 0.322 |
| Among-individual covariance | Response to prey – Web diameter | -0.083 | -0.746 | 0.431 |
| Among-individual covariance | Response to prey – Web density | 0.073 | -0.521 | 0.809 |
| Among-individual covariance | Response to prey – Web symmetry | -0.001 | -0.164 | 0.167 |
| Among-individual covariance | Response to prey – Body length | -0.107 | -0.961 | 0.581 |
| Among-individual variance | Reaction to predation threat | 0.292 | 0.00000 | 0.672 |
| Among-individual covariance | Reaction to predation threat – Web diameter | 0.170 | -0.075 | 0.440 |
| Among-individual covariance | Reaction to predation threat – Web density | -0.192 | -0.470 | 0.085 |
| Among-individual covariance | Reaction to predation threat – Web symmetry | -0.025 | -0.154 | 0.079 |
| Among-individual covariance | Reaction to predation threat – Body length | 0.279 | -0.011 | 0.620 |
| Among-individual variance | Web diameter | 0.507 | 0.220 | 0.816 |
| Among-individual covariance | Web diameter – Web density | -0.340 | -0.615 | -0.084 |
| Among-individual covariance | Web diameter – Web symmetry | 0.009 | -0.120 | 0.143 |
| Among-individual covariance | Web diameter – Body length | 0.359 | 0.088 | 0.657 |
| Among-individual variance | Web density | 0.609 | 0.255 | 0.976 |
| Among-individual covariance | Web density – Web symmetry | 0.014 | -0.134 | 0.179 |
| Among-individual covariance | Web density – Body length | -0.460 | -0.778 | -0.165 |
| Among-individual variance | Web symmetry | 0.058 | 0 | 0.175 |
| Among-individual covariance | Web symmetry – Body length | 0.044 | -0.130 | 0.224 |
| Among-individual variance | Body length | 1.079 | 0.683 | 1.527 |
| Among-date variance | Response to prey | 73.338 | 30.782 | 126.807 |
| Among-date variance | Reaction to predation threat | 1.207 | 0.488 | 2.171 |
| Among-date variance | Web diameter | 0.064 | 0.016 | 0.139 |
| Among-date variance | Web density | 0.134 | 0.038 | 0.271 |
| Among-date variance | Web symmetry | 0.083 | 0.016 | 0.181 |
| Among-date variance | Body length | 0.00001 | 0.00001 | 0.00001 |
| Residual variance | Response to prey | 45.674 | 36.237 | 55.645 |
| Residual covariance | Response to prey – Reaction to predation threat | -0.320 | -1.696 | 1.070 |
| Residual covariance | Response to prey – Web diameter | -0.040 | -0.813 | 0.713 |
| Residual covariance | Response to prey – Web density | -0.023 | -0.728 | 0.665 |
| Residual covariance | Response to prey – Web symmetry | -0.103 | -1.115 | 0.903 |
| Residual covariance | Response to prey – Body length | -0.00003 | -0.009 | 0.009 |
| Residual variance | Reaction to predation threat | 1.837 | 1.391 | 2.326 |
| Residual covariance | Reaction to predation threat – Web diameter | -0.013 | -0.187 | 0.180 |
| Residual covariance | Reaction to predation threat – Web density | 0.107 | -0.070 | 0.279 |
| Residual covariance | Reaction to predation threat – Web symmetry | 0.113 | -0.120 | 0.352 |
| Residual covariance | Reaction to predation threat – Body length | 0.00001 | -0.003 | 0.003 |
| Residual variance | Web diameter | 0.579 | 0.451 | 0.733 |
| Residual covariance | Web diameter – Web density | -0.144 | -0.240 | -0.053 |
| Residual covariance | Web diameter – Web symmetry | 0.173 | 0.052 | 0.299 |
| Residual covariance | Web diameter – Body length | -0.00001 | -0.004 | 0.004 |
| Residual variance | Web density | 0.464 | 0.351 | 0.585 |
| Residual covariance | Web density – Web symmetry | -0.185 | -0.303 | -0.072 |
| Residual covariance | Web density – Body length | -0.00000 | -0.003 | 0.004 |
| Residual variance | Web symmetry | 0.951 | 0.748 | 1.161 |
| Residual covariance | Web symmetry – Body length | -0.00003 | -0.006 | 0.006 |
| Residual variance | Body length | 0.0001 | 0.0001 | 0.0001 |
